# Rescued from extinction in two buckets: genomic consequences of an extreme bottleneck in the endangered Owens pupfish

**DOI:** 10.64898/2026.10.26.761965

**Authors:** David Tian, Bruce J. Turner, Christopher H. Martin

## Abstract

Rapid population bottlenecks can threaten population viability by reducing genetic diversity and effective population size while increasing inbreeding. Freshwater fishes are particularly vulnerable to bottlenecks due to habitat fragmentation, water extraction, and invasive species. The Owens pupfish (*Cyprinodon radiosus*) is an endangered freshwater fish endemic to Owens Valley, CA that was originally thought to be extinct prior to rediscovery in 1964. In 1969, Edwin (Phil) Pister famously saved the entire species from extinction in two buckets after learning that the sole spring they inhabited was drying up. Today, six extant refuge populations ranging in size from 100 - 10,000 individuals remain, which all descend from this severe bottleneck of an estimated 400 individuals. Here, we sequenced whole genomes of 12 individuals from two populations (one extirpated, one extant) to reconstruct recent demographic history and examine the genomic consequences of this well-documented bottleneck on genetic diversity, effective population size, and inbreeding in the Owens pupfish. We estimated the timing of the bottleneck precisely to 1977-1985, closely coinciding with its documented history. Genetic diversity is low across the genome and the bottleneck caused a 17-fold contraction in effective population size over the span of 8 generations. We found low levels of inbreeding that likely predated the bottleneck. Our results suggest that Owens pupfish are threatened by reduced genetic diversity and low effective population size which may constrain adaptive potential, but that recent inbreeding is unlikely to be a serious concern due to population expansion. Our study highlights how the long-term effects of bottlenecks depend not only on severity, but also duration and the rate of population growth afterwards.

## Introduction

Populations that have undergone extreme bottlenecks are vulnerable to demographic stochasticity and genetic threats of reduced genetic diversity and increased inbreeding and drift, which can limit adaptive capacity and exacerbate extinction risk (Lande 1988; Melbourne and Hastings 2008; Willi et al. 2022). However, the long-term genomic consequences of a bottleneck depend on how sharply populations decline, the minimum size reached, how quickly population expansion occurs, and chance (Kirkpatrick and Jarne 2000; Bouzat 2010). Consequently, it is crucial to assess key conservation genetic parameters such as effective population size, genetic diversity, and inbreeding to determine the effects of population decline for a given species and subsequent management options (Hoban et al. 2021). These assessments allow us to infer the adaptive potential of populations, their susceptibility to drift, and whether they may face reduced fitness due to inbreeding depression (Mable 2019), all of which are essential for developing targeted conservation management plans that can monitor and guide populations before local extinction occurs (Rossetto et al. 2021).

Conservation issues of population decline are especially pertinent in freshwater ecosystems and fishes due to anthropogenic factors such as habitat alteration, water extraction, and non-native species that pose threats such as competition and predation (Jelks et al. 2008). Since 1970, freshwater populations have declined by 83% (Harrison et al. 2018) and the modern extinction rate for North American freshwater fishes is 877X greater than the background rate (Burkhead 2012). Among freshwater fishes, the American Southwest in particular has high levels of endemism due to isolated arid environmental conditions and high rates of extinction, with many species comprising a series of fragmented and relic populations (Olden and Poff 2005).

A prime example is the Owens pupfish (*Cyprinodon radiosus*) (Miller 1948), which can serve as a model for understanding bottleneck effects due to its well-known recent demography and near-extirpation. Historically, the species was reported throughout the Owens Valley region in Inyo County, CA along the Owens River and its tributaries (Kennedy 1916). Once so abundant they were a staple food source for local indigenous communities (Lawton et al. 1976), widespread introduction of non-native game fish and diversion of water from the Owens River to the Los Angeles metropolitan area are believed to have contributed to its decline and extirpation across much of its range, to the point that Owens pupfish were thought to be extinct in 1942 (Miller 1969). However, a single remnant population of ∼200 individuals was re-discovered in 1964 at Fish Slough (Miller and Pister 1971). In 1969, ∼800 individuals from the Fish Slough area were famously rescued from extinction by Phil Pister as the local spring began to rapidly dry (Miller and Pister 1971; Pister 1993). Four translocations were made within the slough area but BLM Spring (∼400 individuals) was the only successful one, and is the source from which all subsequent extant populations are derived (Miller and Pister 1971; Pister 1993). Today, the species persists distributed across populations that have been repeatedly founded, extirpated by human-assisted introductions of invasive predators such as largemouth bass, re-founded, and translocated, with six extant populations (U.S. Fish and Wildlife Service 2022).

Current conservation strategies have thus far primarily focused on protecting and monitoring multiple refuge populations (U.S. Fish and Wildlife Service 2022). While restoring habitat, removing non-native species, and translocations are helpful and important, long-term viability likely also depends on maintaining genetic diversity for adaptive potential (Kardos et al. 2021). A previous study based on nine microsatellite loci indicated that genetic diversity and effective population size were low across extant refuge populations and that some populations were becoming more differentiated over time. However, the extent to which effective population size has declined from historical levels and inbreeding is manifest across the genome as a result of the bottleneck remain unclear. Furthermore, microsatellite-based measures of genetic diversity are more biased and less accurate than genomic measures (Fischer et al. 2017; Zimmerman et al. 2020), which also allow for investigation of how genetic diversity is distributed across the genome.

Here, we leverage a genomic dataset to explore the effects of a severe bottleneck in Owens pupfish by inferring effective population size over the last 150 generations and estimating genome-wide genetic diversity and inbreeding. We sampled a total of 12 genomes from an extirpated population at White Mountain Research Station (WMRS) founded in 1988 and extirpated by 1995, and an extant population at Well 368, a flowing artesian well, which hosts a population of an estimated 100 - 1,000 individuals (U.S. Fish and Wildlife Service 2022). Owens pupfish provide an excellent case study to explore the effects of severe bottlenecks and how genomic perspectives can aid contemporary management strategies in an iconic species of significant conservation and cultural significance.

## Results

### Population structure

Principle component analyses based on 91,894 LD-pruned SNPs from a total of 1,836,309 SNPs suggested the presence of some population structure between WMRS and Well 368, with PC1 explaining 13.5% of variation (Fig. 1). However, admixture analyses based on the same set of LD-pruned SNPs indicated a lack of population structure as K = 1 was the best supported value for the number of population clusters. Genome-wide *F*_ST_ analyses similarly suggested low or minimal population differentiation (Weir and Cockerham’s *F*_ST_ = 0.05; Hudson’s *F*_ST_ = 0).

**Figure 1.**
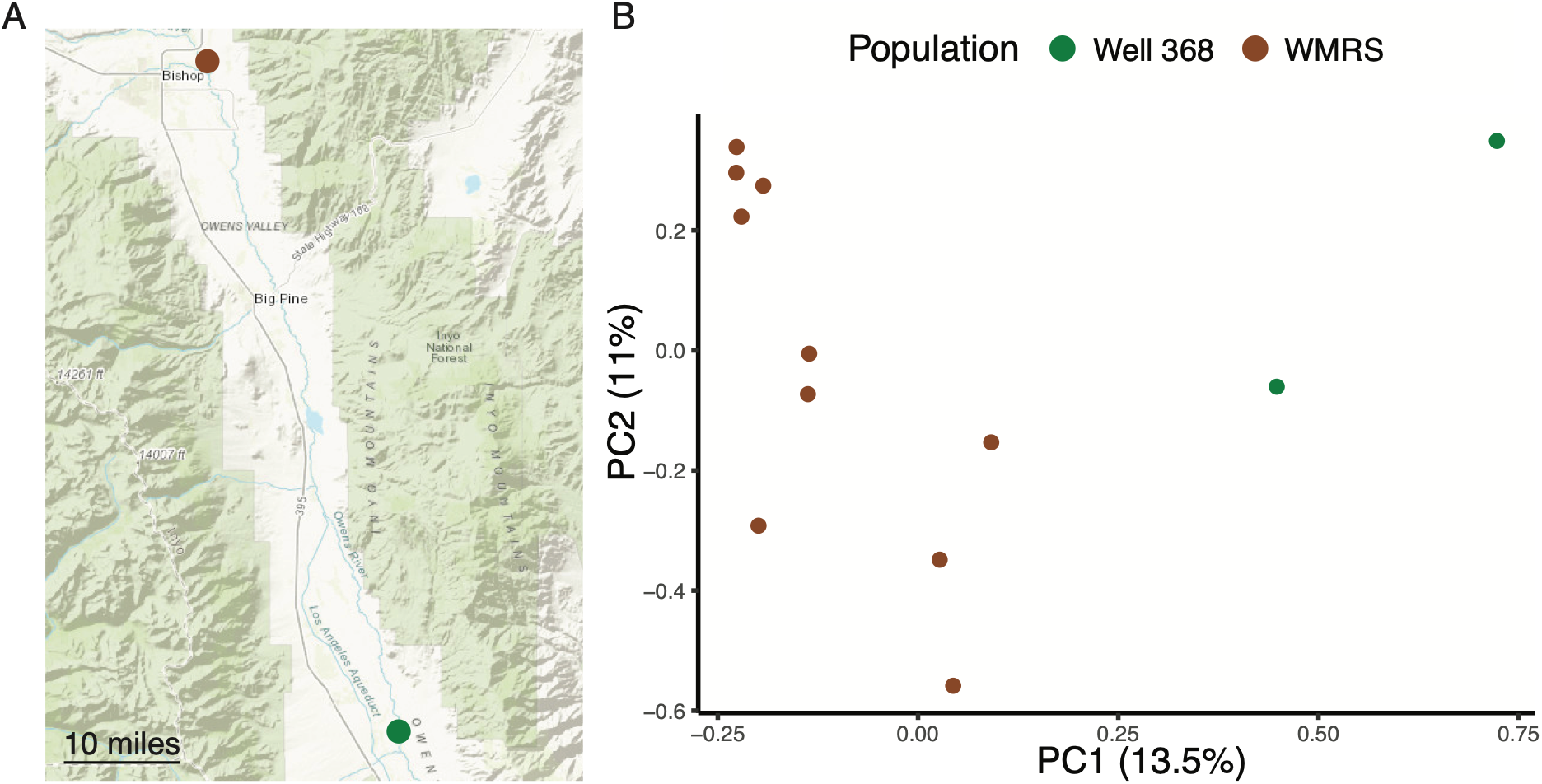
Sampling locations and population structure. A) Map of Owens pupfish sampling locations in Owens Valley, CA with locations of White Mountain Research Station (WMRS) and Well 368 shown in brown and green, respectively. Map adapted from Data Basin. B) Principle component analysis indicating low to moderate population structure.

### Historical and recent demographic history

To infer historical effective population size, we leveraged SMC*++* (Terhorst et al. 2017), a coalescent approach for estimating demographic history from multiple unphased genomes. We found a sharp reduction in effective population size from ∼30,000 to ∼5,000 roughly 10 – 20 kya, consistent with when the Owens Valley began to dry (Bacon et al. 2006) (Fig. 2A). While coalescent approaches can infer deeper demographic history well, they lack resolution on more recent timescales. Consequently, we also inferred recent effective population size over the past 150 generations using GONE2, a linkage-disequilibrium based approach (Santiago et al. 2025).

**Figure 2.**
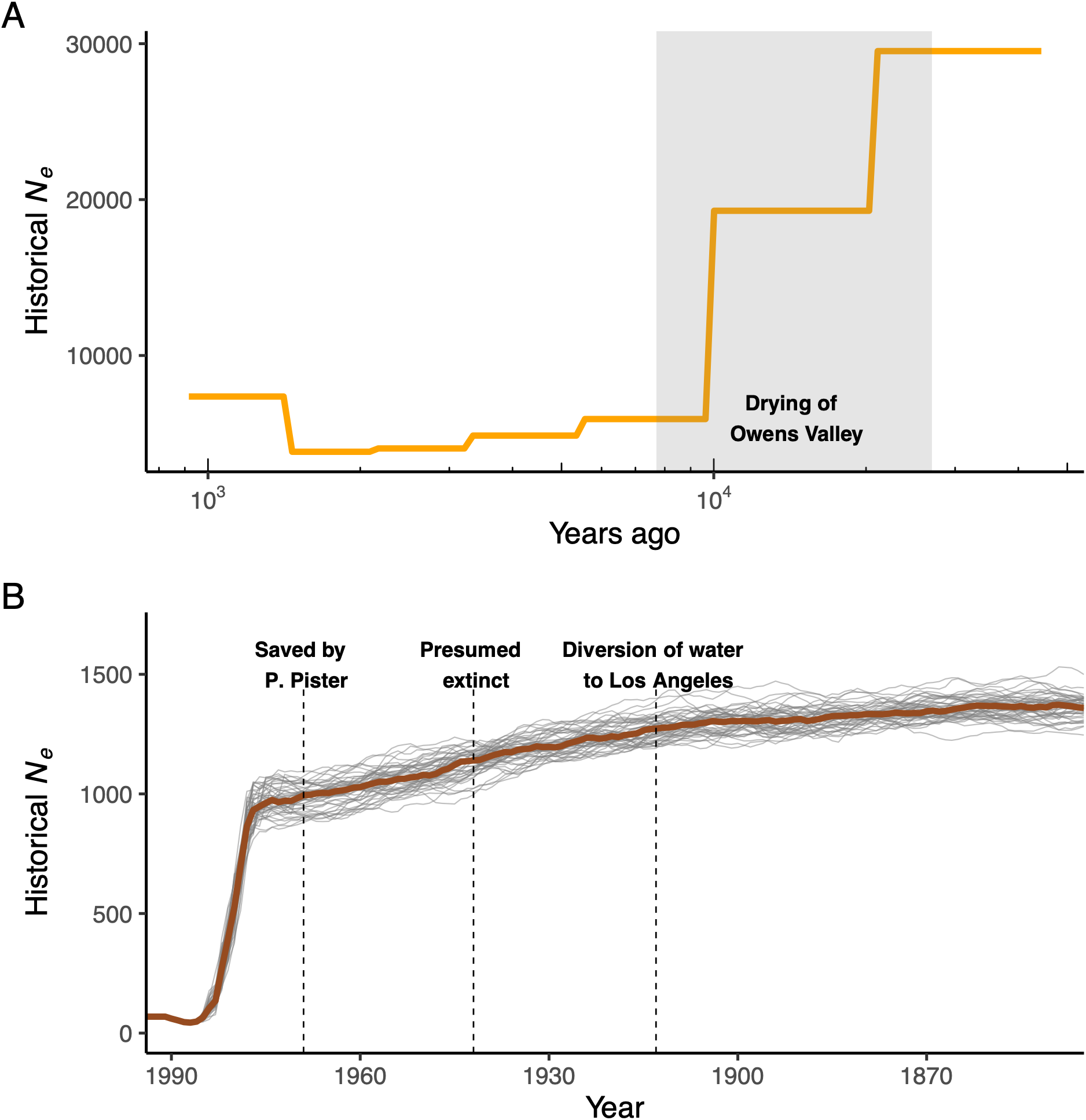
Historical and recent demographic trajectories of Owens pupfish. A) Historical effective population size estimated with SMC*++* (Terhorst et al. 2017). Shaded area denotes late Pleistocene when the upper Owens Valley began to dry (7,700 – 27,000 ya) based on stratigraphic data and radiocarbon dating from Bacon et al. (2006). B) Recent effective population size over the past 150 generations estimated with GONE2 (Santiago et al. 2025) based on 10 individuals from the White Mountain Research Station population. Here, we assume that generation time is one year. Grey lines represent 50 bootstraps with the brown line representing the median estimate. Dashed vertical lines mark when water was diverted from the Owens River to Los Angeles (1913), when Owens pupfish were presumed extinct (1948), and when P. Pister saved the species in two buckets (1969).

We found that effective population size was relatively stable at around 1,250 before a gradual decline in the early 20^th^ century down to about 1,000, coinciding with when stocking of game fish and diversion of water from the Owens River to Los Angeles first began (Fig. 2B). While the timing and severity of the contemporary bottleneck in Owens pupfish are well documented (Pister 1993; Finger et al. 2013), it remains unclear how extensive the reduction in effective population size was. We found a sharp reduction in effective population size that closely coincided with the known bottleneck of 1969, where effective population size remarkably dropped by a factor of over 17X from 831 – 48 in the span of just 8 generations (1977 – 1985) (Fig. 2B). These results were slightly influenced by the recombination rate used, with higher recombination rates leading to lower estimates of effective population size prior to the bottleneck (Supp. Fig. 1). However, neither the inferred timing of the bottleneck nor the resulting effective population size were affected.

We estimated contemporary effective population size (1994) of the WMRS population to be 84 (90% CI: 48.07-146.77) using currentNe2 (Santiago et al. 2025), similar to previous estimates of other populations (Finger et al. 2013). With only two individuals for Well 368 we were unable to estimate contemporary effective population size for that population. Consistent with the expectation of strong drift due to the extreme reduction in effective population size as a result of the bottleneck, we found that genome-wide Tajima’s D was close to 0 at 0.26 and 99.8% of windows were between our significance thresholds of -2 and 2, suggesting that the vast majority of the genome is evolving neutrally, consistent with the effects of the bottleneck and small population size (Fig. 3c).

**Fig. 3.**
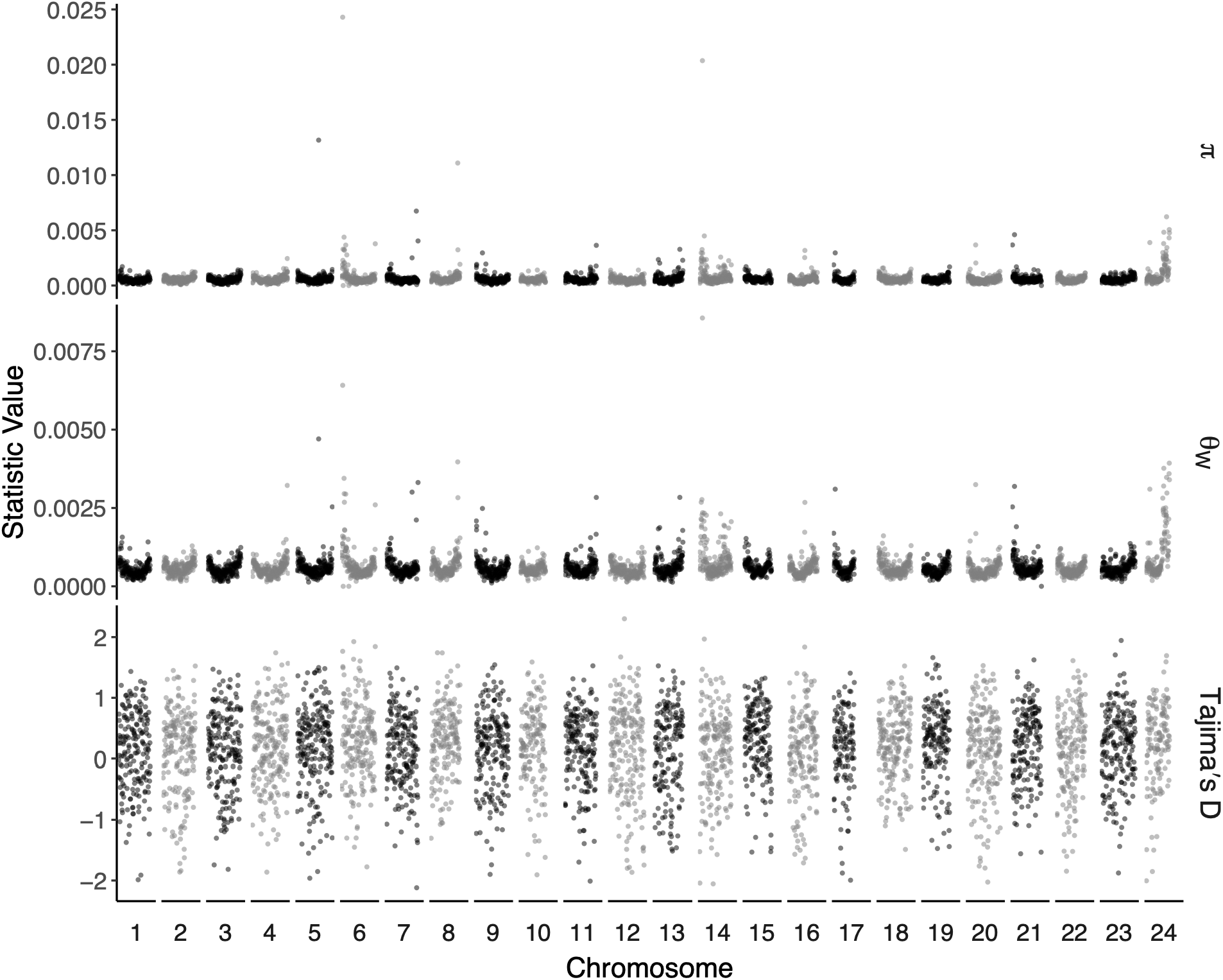
Genome-wide genetic diversity and Tajima’s D. Genome-wide nucleotide diversity, Watterson’s theta, and Tajima’s D in Owens pupfish in non-overlapping sliding windows of 250 kb across all 24 chromosomes. Population genetic statistics were calculated with pixy (Korunes and Samuk 2021).

**Fig. 4.**
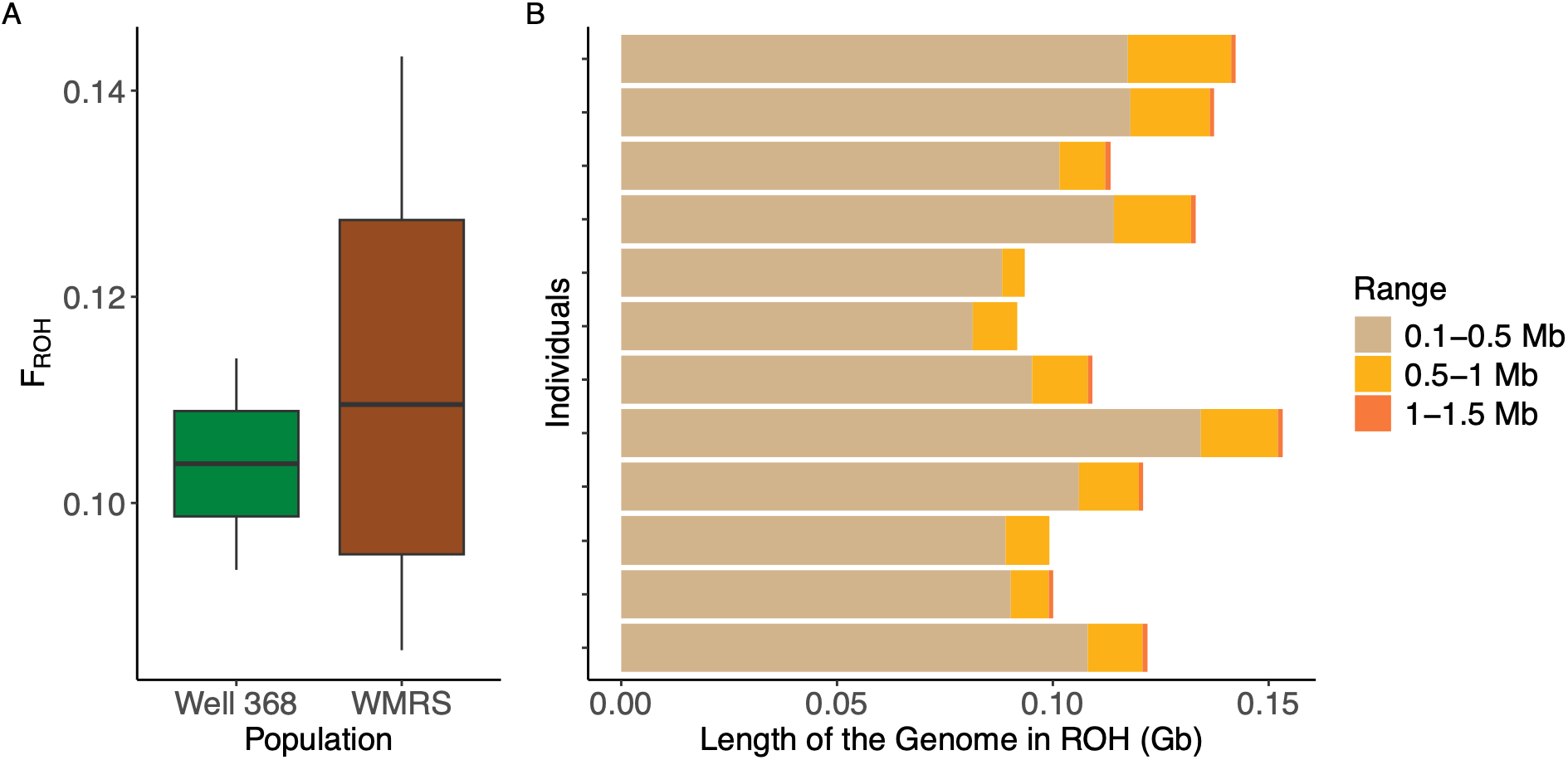
Inbreeding in the Owens pupfish. A) Comparison of the percent of the genome made up of runs of homozygosity greater than 100 kb (F_ROH_) in Well 368 and White Mountain Research Station (WMRS) populations. B) Size distribution of runs of homozygosity per individual.

### Genetic diversity

We found overall low genome-wide nucleotide diversity within Owens pupfish (π = 5.6 x 10^-4^), consistent with their long-term small population size following the drying of Owens Valley during the late Pleistocene (Fig. 2a) and the fact that all existing individuals today are descended from a bottleneck of 400 individuals. Genome-wide Watterson’s theta was similarly low (θ = 5.7 x 10^-4^). While the majority of the genome exhibited consistently low genetic diversity, we found spurious regions of the genome with elevated genetic diversity (Fig. 3). We took an outlier approach and identified genes located in the upper 99^th^ percentile of windows with elevated nucleotide diversity (Supp. Table 1). Gene ontology analyses did not yield significant enrichment of any biological process categories. The elevated genetic diversity towards the end of chromosome 24 may be due to an elevated proportion of repeats (Supp. Fig. 2).

### Inbreeding

In addition to low genetic diversity and effective population size, we found moderate inbreeding within both populations, calculated as the percent of the genome made up of runs of homozygosity (ROHs) at least 100 kb in length (F_ROH_). Overall, F_ROH_ was on average (0.11), with non-significant differences between the WMRS and Well 368 populations (WMRS F_ROH_ = 0.11, Well 368 F_ROH_ = 0.10; *P* = 0.58, T-test). The vast majority (88.2%) of ROHs per individual were 100 – 500 kb in length. The average size across individuals was 206 kb, suggestive of historical inbreeding dating ∼87 generations back to the early 20^th^ century, rather than recent inbreeding as a result of the bottleneck and subsequent founding of populations.

## Discussion

Our study provides the first genomic assessment of Owens pupfish and demonstrates how the effects of bottlenecks depend not only on severity but also duration. Our findings suggest large declines in effective population size associated with the drying of Owens Valley and the 1969 bottleneck when the species was miraculously saved from extinction. Consistent with low effective population size, we found low genetic diversity. However, we also found that inbreeding has remained at low to moderate levels that can primarily be traced back to historical rather than recent close matings. While microsatellite-based approaches have the advantage of more easily sampling large numbers of individuals and populations, here we highlight how even a limited number of whole genomes can provide deep and relevant insights into questions of conservation concern.

### Recent demographic history

Based on patterns of genome-wide linkage-disequilibrium, we inferred an initial decline in effective population size that coincides with early 20^th^ century human-associated ecological and habitat intervention, followed by an extreme reduction in effective population size from 831 – 48 from 1977 to 1985. This single event relegated Owens pupfish to a reduced effective population size below the recommended minimum viable population size of 500 for maintaining evolutionary potential (Frankham et al. 2014). Although the bottleneck occurred in 1969, effective population size is estimated to have remained consistent in the 800s between 1969 - 1977. This lag time may be due to error in the recombination rate used, overlapping generations, or large numbers of offspring reaching reproductive maturity, thus evening out or reducing variance in reproductive success (Gargiulo et al. 2025). Similar lags in effective population decline relative to census demographic decline have been observed in Delta Smelt (Finger et al. 2026). Our estimate of effective population size in the extirpated WMRS population of 84 in 1994 was consistent with previous microsatellite-based estimates of effective population size in other populations (BLM Spring, Marvin’s Marsh, Letter Ponds, Well 368, Lower Mule Spring) from 2007-2011 (Finger et al. 2013).

### Population differentiation

Previous work found generally low but significant pairwise *F*_ST_ values ranging from 0.03 – 0.100 between many populations. More specifically, *F*_ST_ between Well 368 in 2010 and other populations ranged from 0.067 – 0.089 (Finger et al. 2013). Our genome-wide Weir and Cockerham’s estimate of *F*_ST_ (Weir and Cockerham 1984) between WMRS (1994) and Well 368 (2010) was very similar (0.05), suggesting congruence in previous microsatellite and current SNP based estimates. However, our Hudson’s estimate of *F*_ST_ (Hudson et al. 1992) was zero, indicating no population differentiation. Given differences in sample size between WMRS and Well 368, Hudson’s *F*_ST_ is preferred as it is independent of sample size and does not systematically overestimate *F*_ST_, while Weir and Cockerham’s *F*_ST_ is dependent on the ratio of samples sizes (Bhatia et al. 2013). Furthermore, microsatellite-based estimate of *F*_ST_ are often overestimated relative to genome-wide SNPs, due to orders of magnitude fewer markers, ascertainment bias, and higher sequence variance in microsatellite regions (Fischer et al. 2017). As a result, overall divergence among populations of Owens pupfish may be lower than previously thought, although we caution that we were able to compare only two populations.

### Genetic diversity

Genome-wide estimates and distributions of nucleotide diversity and Watterson’s theta indicated that Owens pupfish harbor low genetic diversity, consistent with previous estimates of low heterozygosity relative to other desert pupfishes, with the exception of the extremely isolated Devils Hole pupfish (Tian et al. 2026). The uniform distribution of genetic diversity across the genome suggests that effective population size has likely been consistently small for an extended period of time, consistent with our demographic history analyses. A few regions of locally elevated genetic diversity were observed which may reflect regions under balancing selection or regions of elevated local recombination. However, some of these hot spots maybe be due to elevated levels of repeat content. Overall, the low genetic diversity of Owens pupfish suggests that adaptive potential to anthropogenic selective pressures may be limited (Kardos et al. 2021).

### Inbreeding

A potential reason why inbreeding may not be extensive is the propensity for rapid population growth in Owens pupfish, so long as suitable habitat is available. Following the bottleneck, 400 remnant individuals were transplanted to BLM Spring in August 1969 and less than a year later, the population was conservatively estimated to be at least 1,000 individuals (Miller and Pister 1971). In another example, in April 1977, 300 individuals were introduced into an impounded section of BLM Spring. By September 1977, the population had expanded to 1,925 (CI: 975-3,915) and a year later, up to 9,585 (CI: 6,500-13,000) (Young 1979). Furthermore, Mire and Millet (1994) found that female Owens pupfish spawn up to 200 times daily, with 38% of spawning acts resulting in egg deposition, 95% of which are fertilized, suggesting high fecundity.

While effective population size did not recover following the bottleneck, census population size rapidly did, potentially allowing for the avoidance of inbreeding. The overall small size of ROHs among our samples suggest that the effects of serial translocations and founder events following the bottleneck have not resulted in significant inbreeding. Based on overall low genetic diversity and few long ROHs (> 1 Mb), we infer Owens pupfish to have low risk of inbreeding depression based on the IDrisk metric (Kyriazis et al. 2025). While the reason behind the extirpation of the WMRS population in 1995 is unclear, inbreeding depression is unlikely to have been the primary cause.

### Future conservation

When the Owens pupfish was first described, it was thought to be extinct (Miller 1948) and in 1969, the Owens pupfish nearly did go extinct as the sole spring it inhabited at Fish Slough began to dry (Pister 1993). The fact that this species continues to persist despite large reductions in effective population size and low genetic diversity is likely due to rapid population expansion following the bottleneck along with careful monitoring and protection of habitat. Future sampling of historical specimens will allow us to test whether purging may have played a role in the post-bottleneck rebound of Owens pupfish.

In 2021, the most recent sixth population of Owens pupfish was founded at River Spring Lakes Ecological Reserve, sourced from all five other extant populations (U.S. Fish and Wildlife Service 2022). Moving forward, integrating genomic sampling across all populations will be essential for improving the long-term viability of Owens pupfish to guide translocations and further assess how genetic diversity, inbreeding, and population differentiation may change over time.

## Materials and Methods

### Sample acquisition

In total, we sequenced 12 new whole genomes of *Cyprinodon radiosus*. 10 individuals were sampled from White Mountain Research Station in 1994, while 2 individuals were sampled from Well 368 in 2010. We extracted DNA from fin clips for adults with the Qiagen Dneasy Blood & Tissue kit. Library preparation and sequencing was performed by the QB3-Berkeley Genomics core labs (QB3 Genomics, UC Berkeley, Berkeley, CA, RRID:SCR_022170). DNA quality was assessed with an Agilent Fragment Analyzer and libraries were prepped using the KAPA HyperPrep kit for DNA. Libraries were sequenced on an Illumina NovaSeq 6000 150PE S4 flowcell, generating 20 Gb minimum of data per sample. All sample metadata may be found in Supp. Table 2.

### Genotyping and variant calling

Reads were trimmed with fastp v0.23.4 (Chen et al. 2018) and mapped to the UCB_CyNevMio_1.0 *Cyprindon nevadensis mionectes* reference genome (GCA_030533455.1) with BWA mem v0.7.17 (Li 2013) using default parameters. Duplicates were marked with MarkDuplicatesSpark from GATK v4.5.0.0 and coverage and mapping quality for each sample was assessed with qualimap v2.2.2 (Okonechnikov et al. 2016). We called variants for each individual with GATK Haplotype Caller (-ERC GVCF) v4.5.0.0 (McKenna et al. 2010) and conducted joint genotyping with GATK GenotypeGVCFs to generate an allsites VCF. Variant sites and invariant sites were separated, and missing genotypes were reset to ./. from 0/0 due to how GATK represents missing genotypes in version 4.5.0.0 (Caetano-Anolles 2024). A genmap mappability mask (-k 150 -E 2) for regions of the genome that uniquely map (mappability > 0.5) was applied. Variant sites were filtered with GATK recommended hard filters (QD < 2, QUAL < 30, SOR > 3, FS > 60, MQ < 40, MQRankSum < -12.5, ReadPosRankSum < -8) and for biallelic SNPs (DePristo et al. 2011; Van der Auwera et al. 2013). Furthermore, we filtered SNPs to have minimum FMT/GQ of 20, a minimum read depth FMT/DP of 3, and site-level depth INFO/DP of between 1/3X and 2X the mean for minimums and maximums, respectively. We removed 88 SNPs where ALT = *. One sample (CYPRAD18_1) was removed due to high individual missingness (> 40%), calculated by vcftools (v0.1.16) (Danecek et al. 2011). In the end, we ended up with a final set of 1,836,309 SNPs. Invariant sites were filtered to have RGQ >= 20 and FMT/DP >= 3. Filtered variant and invariant VCFs were merged as input for pixy to provide unbiased estimates of population genetic statistics (see below).

### Historical and recent demography

We ran SMC*++* to infer historical effective population size trajectories and varied the chosen distinguished individuals across all 12 individuals to generate distinct datasets for estimation and a composite-likelihood (Terhorst et al. 2017). To infer more recent demographic trajectories, we used GONE2 to infer effective population size over the last 150 generations using the WMRS population (n = 10) (Santiago et al. 2025). We assumed a recombination rate of 2.8 cM/Mb based on the median value among fishes (Stapley et al. 2017) and ran GONE2 at varying lower and higher rates (2.3 and 3.3 cM/Mb) to test the effect of recombination rate (Supp. Fig. 1). Finally, we leveraged currentNe2, a linkage-disequilibrium based method to also estimate contemporary effective population size based on integration over the whole genome (Santiago et al. 2025).

### Population structure

We pruned SNPs in strong linkage disequilibrium using the pruning function (--indep-pairwise 50 10 0.2 in PLINK (v1.9) (Chang et al. 2015), which left 91,894 SNPs. We then characterized population structure following two complementary approaches. First, we used PLINK to conduct principal component analysis using the –pca function. Afterwards, we used ADMIXTURE (v1.3.0) (Alexander et al. 2009) to assign individuals to varying numbers of population clusters (K = 1-5). Based on cross-validations errors, K = 1 was the value that best assigned individuals to population clusters. Finally, we also assessed population differentiation in terms of *F*_ST_ using pixy (v2.2.1) (Korunes and Samuk 2021). We calculated Weir and Cockerham’s *F*_ST_ (Weir and Cockerham 1984) for ease of comparison to previous studies and Hudson’s *F*_ST_ (Hudson et al. 1992), which is more robust to unequal sample sizes (Bhatia et al. 2013).

### Genetic diversity and Tajima’s D

We measured genetic diversity across our entire dataset in terms of π, the average number of pairwise nucleotide differences between all possible pairs of individuals in a population, and Watterson’s θ, which quantifies the number of polymorphic sites, with pixy (v2.2.1) (Korunes and Samuk 2021). We ran pixy in sliding windows of 250 kb and calculated genome-wide average π as the sum of the differences divided by the sum of the comparisons across windows. We identified outlier windows of elevated π based on windows that had a π value in the 99^th^ percentile or above (π > 2.5 x 10^-3^) and intersected these outlier windows with our annotation gff file using bedtools v2.30.0 (Quinlan and Hall 2010). We inputted these outlier genes into ShinyGO v0.85 (Ge et al. 2020) and PANTHER v19.0 (Thomas et al. 2022) to test for significant gene ontology categories.

We calculated Watterson’s theta as the sum of per-site theta values across windows divided by the total number of sites with at least one valid genotype. Finally, we used pixy to estimate Tajima’s D based on the difference in mean number of pairwise differences and number of segregating sites in sliding windows of 250 kb, to infer whether certain regions of the genome were evolving neutrally or not.

### Inbreeding

We identified runs of homozygosity (ROHs) in our Owens pupfish samples using the BCFtools/ROH command with the Genotypes option (v1.16) (Narasimhan et al. 2016). We calculated individual inbreeding coefficients in terms of F_ROH_, the proportion of the genome made up of ROHs which we calculated as the sum of ROHs greater than 100 kb divided by the sum of the lengths of the 24 autosomes (1,069,364,557 bp). We compared levels of inbreeding between our two Owens pupfish populations with two-sample t-tests. We also classified ROHs into various lengths of 0.1 – 0.5 Mb, 0.5 – 1 Mb, > 1 Mb, and calculated the proportion of the genome that these classes of ROHs comprised in each individual. The age of ROHs were estimated as g = 100 / 2*ROH length (cM), where g is the number of generations to the most recent common ancestor (Thompson 2013). We assumed a generation time of 1 year and an average recombination rate of 2.8 cM/Mb based on the median rate among fishes (Stapley et al. 2017).

## Data availability

Sequence data is available at NCBI’s SRA database under PRJNA1543672. The reference genome used for alignment is available on NCBI at GCA_030533455.1. Code for all analyses is available on the lead author’s github at https://github.com/tiandavid/radiosus.

## Acknowledgements

This research was funded by NSF CAREER 1749764, NIH 5R01DE027052-02, and Hellman Foundation awards to CHM. All research procedures and animal care protocols (AUP-2021-02-14062-1 and AUP-2021-07-14515) were approved by the University of California, Berkeley Animal Care and Use committee. D.T. was supported by an NSF GRFP (DGE 1752814), the Museum of Vertebrate Zoology, and a Philomathia Graduate Fellowship in the Environmental Sciences. We thank Phil Pister for saving the Owens pupfish from extinction.

## Author Contributions

D.T. conceptualized the project. D.T. performed all analyses and lab work. C.H.M. provided funding for sequencing and analysis and acquired samples. D.T. wrote the manuscript. B.T. collected samples from White Mountain Research Station. All authors edited and reviewed the manuscript.

