## Supplemental Materials for "Rescued from extinction in two buckets: genomic consequences of an extreme bottleneck in the endangered Owens pupfish"

**This file includes:**

Supplementary Figures S1 - S2

Supplementary Tables S1 - S2

**Supplemental Figures**

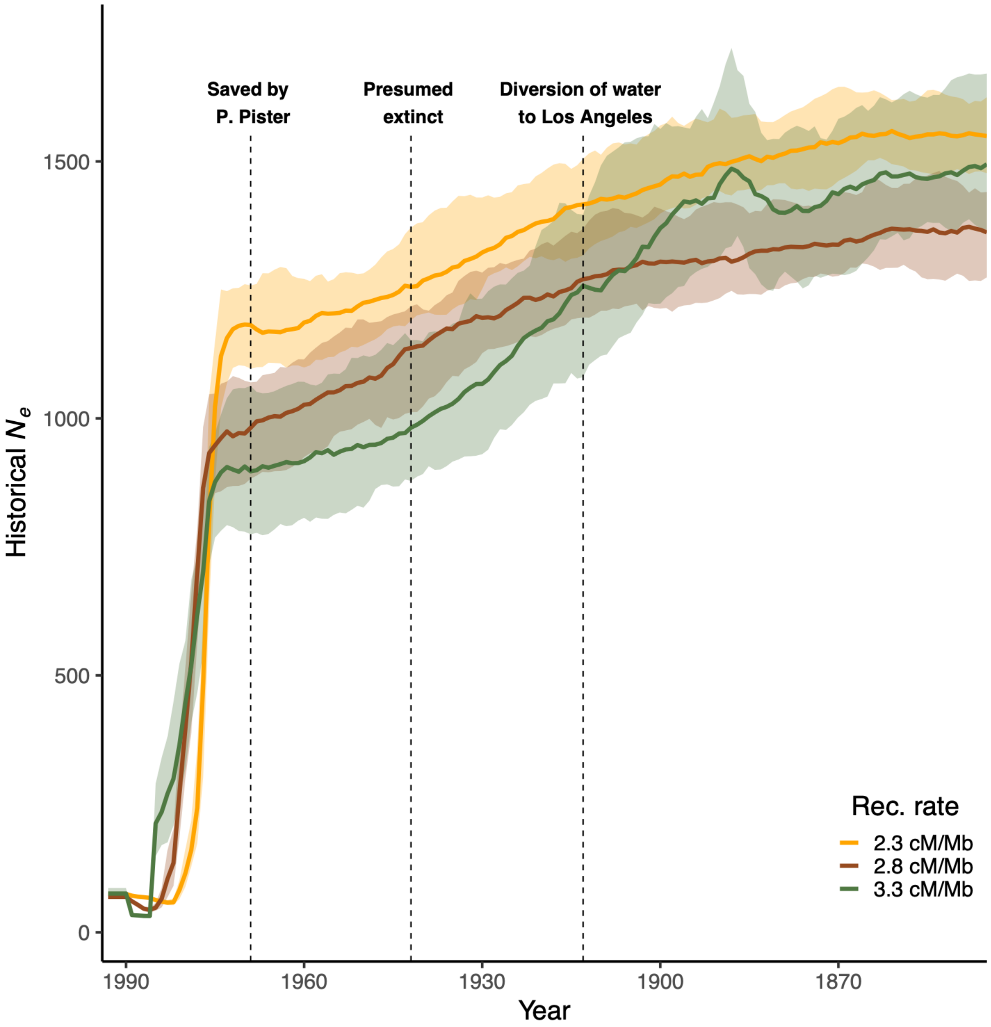


**Supp. Fig. 1.** Recent effective population size of Owens pupfish over the past 150 generations estimated with GONE2 (Santiago et al. 2025) based on 10 individuals from the White Mountain Research Station population. Here, we assume that generation time is one year. Colors represent varying recombination rates used to parameterize GONE2 and shaded areas represent the 95% confidence interval across 50 bootstraps. Dashed vertical lines mark when water was diverted from the Owens River to Los Angeles (1913), when Owens pupfish were presumed extinct (1948), and when P. Pister saved the species in two buckets (1969).

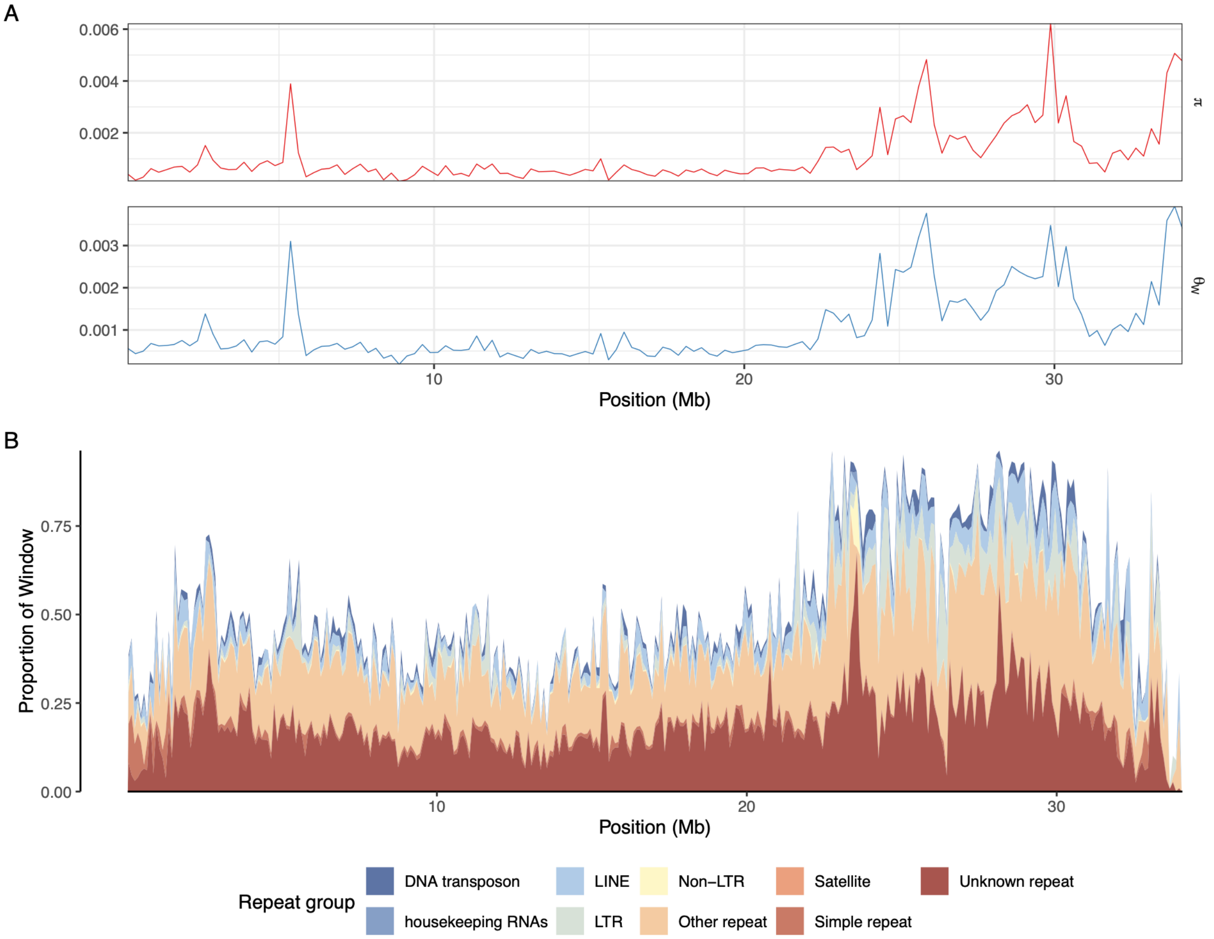


**Supp. Fig. 2.** Genetic diversity along chromosome 24. A) Nucleotide diversity and Watterson's theta in non-overlapping sliding windows of 250 kb calculated with pixy (Korunes and Samuk 2021). B) Distribution of repeat content based on RepeatMasker annotations of the *Cyprinodon nevadensis mionectes* reference genome (GCA_030533455.1).

### Supplemental Tables

**Supp. Table 1.** Known genes found in the upper 99$^{th}$ percentile of nucleotide diversity outlier windows. Gene functions sourced from GeneCards.

| Gene ID | Gene name / function |
|---|---|
| ALOX12B | arachidonate 12-lipoxygenase, 12R type |
| AMICA1 | enables integrin binding activity and protein homodimerization activity |
| CD84 | SLAM family self-ligand receptor and membrane glycoprotein |
| DOCK3 | dedicator of cytokinesis family guanine nucleotide exchange factor |
| Dref | DNA replication-related element factor |
| FCRLA | Fc receptor-like A, an immunoglobulin-like protein related to receptors for the Fc fragment of gamma immunoglobulin G. |
| HAUS7 | subunit of the augmin complex that regulates centrosome and mitotic spindle integrity |
| L1TD1 | LINE-1 Type Transposase Domain-Containing Protein 1 |
| MAP6 | microtubule-associated protein 6 |
| MOG | myelin oligodendrocyte glycoprotein |
| NLRP12 | cytoplasmic NOD-like receptor that attenuates inflammation in activated monocytes |
| NOL9 | nucleolar protein with ATP-dependent polydeoxyribonucleotide 5'-hydroxyl-kinase activity |
| PRKRIR | Protein-Kinase, Interferon-Inducible Double Stranded RNA Dependent Inhibitor |
| SLC25A11 | mitochondrial dicarboxylate carrier that mediates electroneutral exchange of 2-oxoglutarate for malate and other dicarboxylic acids across the inner mitochondrial membrane |
| SPTLC2 | catalytic subunit of serine palmitoyltransferase that initiates sphingolipid biosynthesis |
| TARP | TCR Gamma Alternate Reading Frame Protein |
| TCP11L1 | T-Complex 11 Like 1 |
| TMEM185A | transmembrane protein 185A, a predicted membrane protein that localizes to the CpG island of the fragile site FRAXF |
| TRAV18 | variable region of the T cell receptor alpha chain that contributes to antigen recognition in alpha-beta T cell receptors and supports adaptive immunity |
| TRAV21 | ariable region of the T cell receptor alpha chain that contributes to antigen recognition within peptide–major histocompatibility complexes and helps initiate T cell activation |
| TRIM7 | E3 ubiquitin-protein ligase and member of the tripartite motif (TRIM) family that links selective substrate recognition to ubiquitin-mediated protein degradation |
| ZDHHC24 | ZDHHC palmitoyltransferase 24, a DHHC-type enzyme predicted to catalyze protein-cysteine S-palmitoyltransferase activity |
| ZMYM6 | zinc finger MYM-type protein associated with DNA binding and roles in cytoskeleton organization and regulation of cell morphogenesis |
| ZNF251 | KRAB-domain zinc finger transcription factor that binds DNA and regulates RNA polymerase II-dependent transcription |
| ZNF846 | zinc finger protein 846, a KRAB domain-containing C2H2 zinc-finger transcription factor predicted to bind DNA and regulate RNA polymerase II-dependent transcription |
| wrn-1 | multifunctional RecQ helicase-exonuclease that preserves genomic integrity through ATP-dependent 3'→5' helicase and 3'→5' exonuclease activities |

**Supp. Table 2.** Individual sample metadata. Location abbreviation of WMRS represents White Mountain Research Station. Genome-wide mean coverage and mapping quality were calculated using qualimap (v2.2.2) and individual missingness was calculated with vcftools (v0.1.16).

| Sample ID | Population | Sample Year | Mean coverage | Mapping quality | Ind. missingness |
|---|---|---|---|---|---|
| CRAD_18-10 | WMRS | 1994 | 22.27X | 21.96 | 0.062 |
| CRAD_18-11 | WMRS | 1994 | 24.61X | 22.22 | 0.056 |
| CRAD_18-2 | WMRS | 1994 | 16.96X | 21.73 | 0.083 |
| CRAD_18-3 | WMRS | 1994 | 14.47X | 21.38 | 0.124 |
| CRAD_18-4 | WMRS | 1994 | 18.98X | 21.68 | 0.077 |
| CRAD_18-5 | WMRS | 1994 | 18.90X | 22.19 | 0.078 |
| CRAD_18-6 | WMRS | 1994 | 16.93X | 22.13 | 0.096 |
| CRAD_18-7 | WMRS | 1994 | 13.74X | 21.94 | 0.129 |
| CRAD_18-8 | WMRS | 1994 | 17.27X | 21.90 | 0.082 |
| CRAD_18-9 | WMRS | 1994 | 21.20X | 21.90 | 0.064 |
| CYPRAD18_1 | WMRS | 1994 | 7.18X | 21.59 | 0.458 |
| C_radiosus_W368-2 | Well 368 | 2010 | 12.86X | 21.15 | 0.159 |
| C_radiosus_W368-8 | Well 368 | 2010 | 12.45X | 20.96 | 0.178 |
